# Glucose-Fueled Histone Modifications Drive HIV-1 Latency Reversal at Hypoxia

**DOI:** 10.1101/2025.09.14.676169

**Authors:** Yetunde I. Kayode, Deanna C. Clemmer, Adebola A. Owolabi, AbdulKareem A. AlShaheeb, Elyse K. McMahon, Anna Tangiyan, Lauren Clark, Glenn E. Simmons, Alberto Bosque, Melanie R. McReynolds, Harry E. Taylor

## Abstract

The major barrier to curing HIV-1 infection is the persistence of a latent reservoir in CD4 T cells within tissues which readily fuel viral rebound upon antiretroviral therapy (ART) interruption. Clinical trials aimed at purging these viral reservoirs with latency reversal agents (LRAs) have been unsuccessful owing to our incomplete understanding of the molecular and physiological determinants that underlie latency reversal in these virus-harboring tissues. Here, using a combination of complementary pharmacological and metabolomic approaches, we uncover glucose as a conditionally essential nutrient for HIV-1 latency reversal at hypoxic conditions. By modelling physiological variations in both glucose and oxygen availability as found *in vivo* within tissues that may harbor the HIV reservoir, we show that hyperglycemic conditions potentiate HIV-1 latency reversal. Importantly, we found major classes of clinically relevant LRAs, PKC agonists (PKCags) and histone deacetylase inhibitors (HDACis) have disparate efficacies under glucose-limiting conditions. Mechanistically, we show that this differential glycolytic dependency is due to distinct capacities of LRAs to induce glycolytic flux during adaptation to hypoxia, a condition that increases glycolytic dependence. Furthermore, we show that PKCag-induced glycolysis drives histone lactylation, a post-translational modification (PTM) we found to be associated with HIV-1 latency reversal and promotes increased chromatin accessibility at the HIV promoter. Importantly, we identify KAT2A as a lactyl-transferase critical for histone lactylation induced upon latency reversal. Taken together, our findings uncover glucose and oxygen availability as critical metabolic determinants of HIV-1 latency reversal and underscore the importance of modeling physiologically relevant experimental conditions *in vitro* aimed at identifying therapeutic agents that effectively target the latent reservoir *in vivo*.

## Introduction

Human immunodeficiency virus (HIV) is a CD4 cell tropic retrovirus, which currently infects about 39.9 million people globally^1–3^. The hallmark of HIV infections is a progressive destruction of the primary target of HIV, CD4 T cells, leading to acquired immunodeficiency syndrome (AIDS) in untreated individuals, a condition characterized by unusual susceptibility to opportunistic infections, malignancies and ultimately death^4,5^.

Antiretroviral therapy (ART) has been revolutionizing in the global course of the HIV/AIDS pandemic by prolonging the lives of people living with HIV (PLWH), however, ART has major limitations. ART does not eliminate the latent reservoir of the virus, which is seeded soon after exposure to the virus, hence, viremia rebounds following ART interruption^6–10^. Furthermore, ART does not reconstitute the immune system following initial CD4 T cell depletion, does not alleviate HIV-associated chronic inflammation which causes several co-morbidities including cardiovascular diseases, neurologic disorders, obesity, and is itself associated with toxicity^4,11–15^. Put together, these drawbacks necessitate a functional or sterilizing cure that effectively eliminates the latent reservoir in PLWH, either by reactivation and clearance (latency reversal) or by promoting deep latency. However, the mechanisms that govern HIV latency *in vivo* are not completely understood. The bulk (>98%) of the latent reservoir *in vivo* exist in diverse tissues including the gut, adipose tissues, lymph nodes and the central nervous system^16–21^. Hence, an effective cure must target cellular reservoirs within metabolically diverse anatomical niches. Importantly, many of these tissues are physiologically hypoxic (<5% O_2_), differing in oxygen tensions from the standard laboratory culture conditions (∼21% O_2_) under which therapeutics are developed and screened for^22^. This is important owing to growing evidence that cellular metabolism, which is highly influenced by oxygen availability, is the principal determinant of cellular permissiveness to HIV replication and consequently latency reversal^23–26^. Previous studies aimed at identifying potential biomarkers associated with HIV rebound, have identified predictive metabolic biomarkers following ART interruption and loss of virological control. Specifically, in one study increases in glycolytic end products pyruvate and lactate were shown to be associated with a shorter time to virus rebound following analytical treatment interruption^27^. Similarly, in another report, elite controllers, HIV-infected patients who maintain undetectable viral loads without antiretroviral treatment, displayed increased circulating glycolytic metabolites prior to spontaneous loss of virological control^28^. By contrast, recent studies have demonstrated that downmodulation of glycolysis is a hallmark of cells transitioning into latency^29,30^. Taken together, these studies suggest that targeting HIV and its latent reservoir with pharmacological inhibitors of the glycolytic pathway may be a promising approach toward functional cure and reservoir eradication, but a direct causal and mechanistic link has yet to be established^26,31^.

### Glycolysis is necessary for HIV latency reversal at physiological hypoxia

HIV replication is compromised when glycolysis is inhibited, but HIV latency reversal has not been formally demonstrated to share the same level of sensitivity to glucose restriction in CD4 T cells^24,26,34–37^. Notably, the overwhelming number of *in vitro* HIV replication studies have been executed in cell culture under standard atmospheric oxygen tensions (21% O_2_) which do not mirror the hypoxic (∼1-5% O_2_) microenvironments of the latent reservoir *in vivo*^22,24^. To examine the role of glycolysis in HIV latency reversal under both normoxic and hypoxic conditions, we utilized the J-Lat 5A8 GFP LTR reporter cell line, a CD4 T cell HIV latency model with LRA response properties mimicking primary CD4 T cell models and HIV patient-derived infected cells^38^. Here, we inhibited glycolysis in J-Lat 5A8 cells by pretreating cells for 2 h with 2-deoxyglucose (2-DG), a glucose analog, prior to inducing latency reversal with structurally distinct PKC agonists; PEP005, PMA or bryostatin **(Fig. 1A)**. For these experiments, identical sets of cells were incubated either under standard tissue culture conditions (21% O_2_) or physiologic hypoxia (1% O_2_). We found that inhibition of glycolysis resulted in significant diminution of HIV reactivation, as measured by GFP expression under normoxia (∼50%), but more robust (70-90%) effects were observed under hypoxic conditions for the PKC agonists, thus confirming a role of glycolysis in facilitating latency reversal for this LRA class. To assess whether glycolysis had a universal role in HIV latency reversal rather than a PKC LRA class-specific effect, we also replicated these experiments using structurally distinct HDAC inhibitors romidepsin (RMD), vorinostat (SAHA), or sodium butyrate (BUT) which have been shown to elicit latency reversal via epigenetic mechanisms^39^, a mechanism distinct from PKC agonists **(Fig. 1B)**. Most strikingly, we observed that 2-DG treatment resulted in greater than 95% reduction in latency reversal at hypoxia, thus documenting that glycolysis is necessary for latency reversal by both epigenetic and non-epigenetic LRAs. Moving forward, we performed subsequent experiments with PEP005 and RMD as representative a PKC agonist and HDAC inhibitor, respectively.

**Figure 1.**
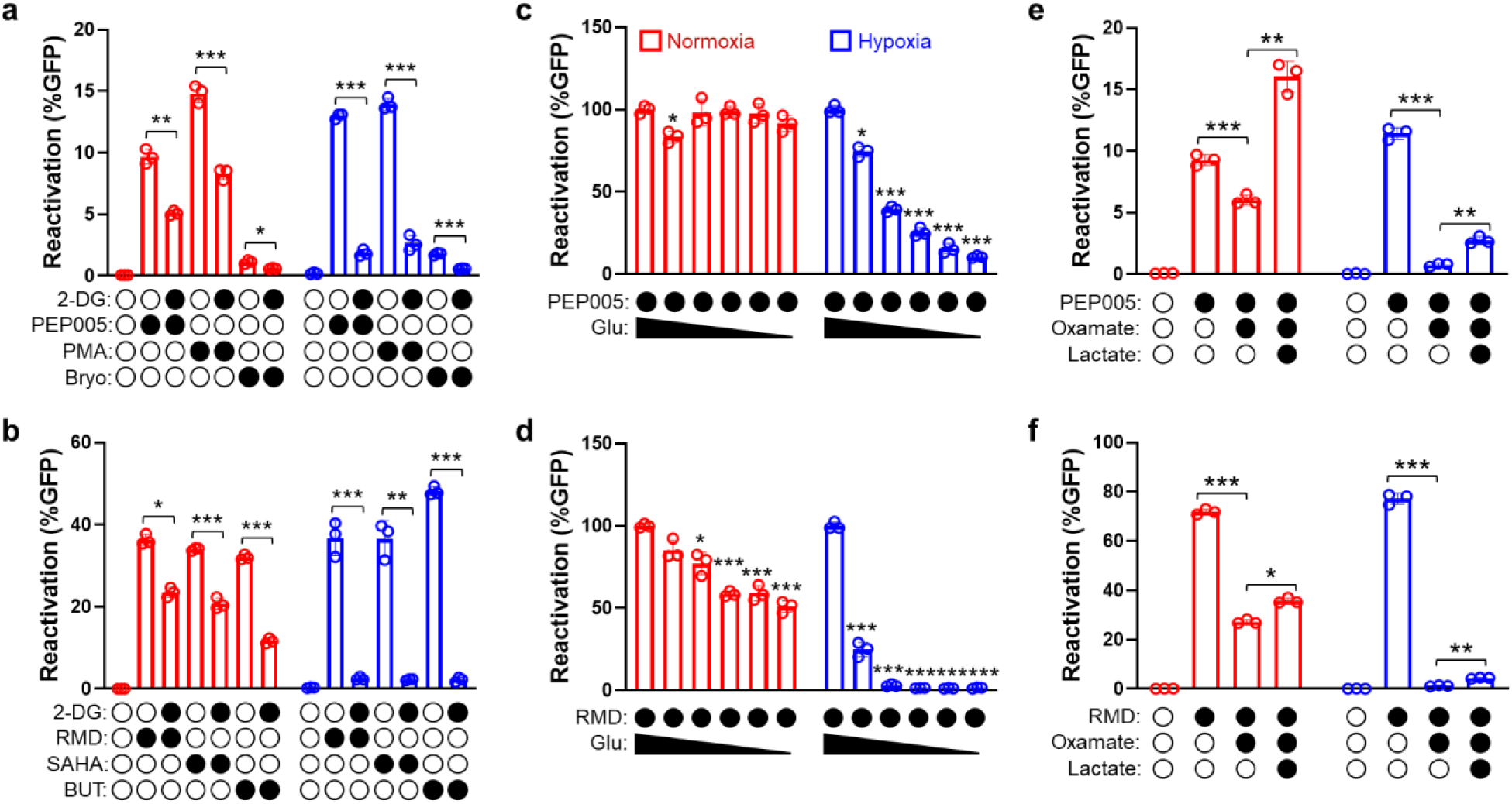
HIV-1 latency reversal is dependent on glucose metabolism. **Glycolytic inhibition blocks HIV latency reversal mediated by multiple LRAs.** Reactivation of latent virus in the J-Lat 5A8 cellular model of HIV latency in cells left untreated or pretreated with glycolytic inhibitor, 2-deoxyglucose, 2-DG (5mM) for 2h under normoxia (21% O_2_) or hypoxia (1% O_2_), then stimulated for 24h with a panel of **(a)** PKC agonists [PEP005 (100nM), PMA (15nM) or bryostatin (100nM)] or **(b)** HDAC inhibitors, RMD (40nM), vorinostat (SAHA, 10mM) or sodium butyrate (BUT, 5mM]. Reactivation of latent virus in J-Lat 5A8 cells Reactivation by **(c)** representative PKC agonist, PEP005 (100nM) and **(d)** representative HDAC inhibitor, romidepsin, RMD (10nM) was measured by GFP fluorescence across media glucose concentrations of 10, 5, 2.5, 1.25 and 0.625mM after 24h of stimulation. Red bars represent samples treated under normoxic conditions (21% O_2_) and blue bars represent samples treated under hypoxic conditions (1% O_2_). Means and SDs of triplicate samples are shown. Statistical significance was determined using paired t-test, comparing reactivation at 10mM glucose to other glucose concentrations in each case. **Exogenous lactate rescues glycolytic inhibition**. Reactivation of latent virus in J-Lat 5A8 cells either untreated or pretreated with glycolytic inhibitor, sodium oxamate (25mM) for 2h under normoxia (21% O_2_) or hypoxia (1% O_2_), then stimulated for 24h with **(e)** PKC agonist, PEP005 (100nM) or **(f)** HDAC inhibitor, RMD (10nM) with or without the addback of exogenous sodium lactate (50mM) during stimulation. Means and SDs of triplicate samples are shown. Statistical significance was determined using paired t-test; *p<0.05, **p<0.005, ***p<0.0005.

### Latency reversal by LRAs demonstrate differential sensitivities to glucose availability

To effectively target the latent reservoir *in vivo*, a LRA must have the capacity to reactivate virus from reservoirs within hypoxic tissues with differential metabolic microenvironments like the gut and lymph nodes^16,21,40^. A key metabolic variable across tissues that has not been fully addressed in the context of HIV latency reversal studies across is glucose availability, as glucose may range from as low as 2.5mM in tissues to ∼5.5mM in venous blood or higher following meals or in diabetic individuals^41^. Our observations define a critical role of glycolysis in HIV latency reversal underscore the importance of understanding the contribution of glucose availability to latency reversal within hypoxic tissue microenvironments. To this end, we tested the impact of glucose levels on HIV latency reversal by performing latency reversal assays while utilizing a glucose titration range that included hypoglycemic (0, 0.625, 1.25, and 2.5 mM), normoglycemic (5 mM), and hyperglycemic (10 mM) levels of glucose under both normoxic and hypoxic conditions. Here, we assessed the capacity of PEP005 and RMD, as representative PKC agonist and HDACi LRAs, respectively, to drive latency reversal over the glucose titration range of 0.625 mM to 10 mM. Here, we observed no glucose concentration-dependent effects on PEP005-induced latency reversal at normoxia. By contrast, executing latency reversal assays with PEP005 under hypoxic conditions uncovered a significant glucose concentration dependence of latency reversal **(Fig. 1C)**. Remarkably, elevated glucose concentrations (10 mM), comparable to those observed in diabetes and to levels present in standard RPMI medium (11.1 mM), significantly enhanced PEP005-induced latency reversal relative to normoglycemic conditions (5 mM). Importantly, unlike latency reversal induced by PEP005, the HDACi RMD demonstrated glucose sensitivity at both normoxia and hypoxia **(Fig. 1D)**. Nonetheless, reactivation by both LRAs under hypoxic conditions was found to be more sensitive to glucose availability than under normoxic conditions. Importantly, this suggests that oxygen levels present within *in vitro* models that do not mirror the hypoxic microenvironment of HIV tissue reservoirs (e.g., the gut and lymph nodes) can mask the physiological efficacy and metabolic dependence of latency reversal agents. This observation highlights the importance of executing studies of latency reversal by utilizing models that recapitulate the metabolic conditions of the tissues that harbor the HIV reservoir.

### Glycolytic end product lactate fuels HIV latency reversal

Hypoxia increases the production of glucose-derived lactate by stimulating glycolysis. This process expands the pool of lactate that is necessary for histone lactylation, a recently discovered epigenetic modification that drives gene expression and effectively couples elevations in cellular glucose-derived lactate to increased gene expression^42^. Interestingly, elevated serum lactate has been associated with increases in HIV expression and virus rebound during analytic treatment interruption (ATI) in HIV patients^27^. We hypothesize that elevated levels of glucose during latency reversal could increase cellular lactate pools that could impact HIV latency reversal. However, it is unknown whether glucose-derived metabolites, like lactate, have a direct role in facilitating latency reversal. To this end, we performed latency reversal-metabolite addback experiments with lactate in either the presence or absence of oxamate, a glycolytic inhibitor that limits the conversion of pyruvate to lactate by lactate dehydrogenase **(Fig. 1E)**. We observed that oxamate-mediated blockade of lactate production and glycolytic flux suppressed latency reversal with PEP005 treatment marginally at normoxia, but potently under hypoxia, thus mirroring the effects of glucose restriction on PEP005-elicited latency reversal **(Fig.1E)**. Importantly, we demonstrated that addition of lactate rescued the effects of oxamate on reactivation. This confirms that the role of glycolysis in latency reversal is mediated, at least in part, by its end product lactate. We obtained comparable results using HDACi RMD as a LRA in experiments **(Fig. 1F)**. To further validate our findings, we took an orthogonal approach to glycolytic flux blockade, by utilizing galactose, a suboptimal glycolytic substrate, as a cell culture carbon source in latency reversal assays^34,43,44^. We observe that HIV latency reversal induced by PEP005 in the presence of galactose was not markedly different than latency reversal in the presence of glucose at normoxia **(Fig. S1)**. By contrast, as observed with glycolytic inhibition with oxamate, hypoxia uncovered significant inhibition of latency reversal through suppression of glycolytic flux by galactose supplementation. Consistent with the effects of oxamate treatment inhibiting latency reversal by reducing the level of lactate production, we were able to rescue the effects of galactose-mediated suppression of glycolysis through the addition of exogenous lactate **(Fig. S1)**. Recent approaches for LRA optimization have involved the combination of different classes of LRAs for a synergistic effect, which is generally achieved using lower doses of each LRA to further minimize toxicity^45^. To evaluate whether glycolysis is also required for the efficacy of LRA combinations, we assessed reactivation using a combination of PEP005 and RMD in the presence of 2-DG. In line with previous observations, reactivation of cells under hypoxia was more significantly sensitive to glycolytic inhibition than cells under normoxia, indicating that the glycolytic requirement for latency reversal is not overcome by synergistic LRA activity, especially under physiologic hypoxia **(Fig. S2).**

Next, we sought to validate our findings in an *in vitro* primary T_CM_ model of HIV latency which utilizes a full-length virus^46^ to rule out contributions of proviral gene deletions^47,48^, elevated baseline glycolysis and dysregulated signaling in the J-Lats^49^. Consistently, we observed a significant reduction of p24 expression in triplicate experiments derived from independent donors even under normoxic conditions, demonstrating that glycolysis is indeed necessary for virus reactivation **(Fig. 2A)**. Taken together, our findings demonstrate that glycolysis is necessary for reactivation of latent HIV and that hypoxic adaptation increases the glycolytic dependence of latency reversal.

**Figure 2.**
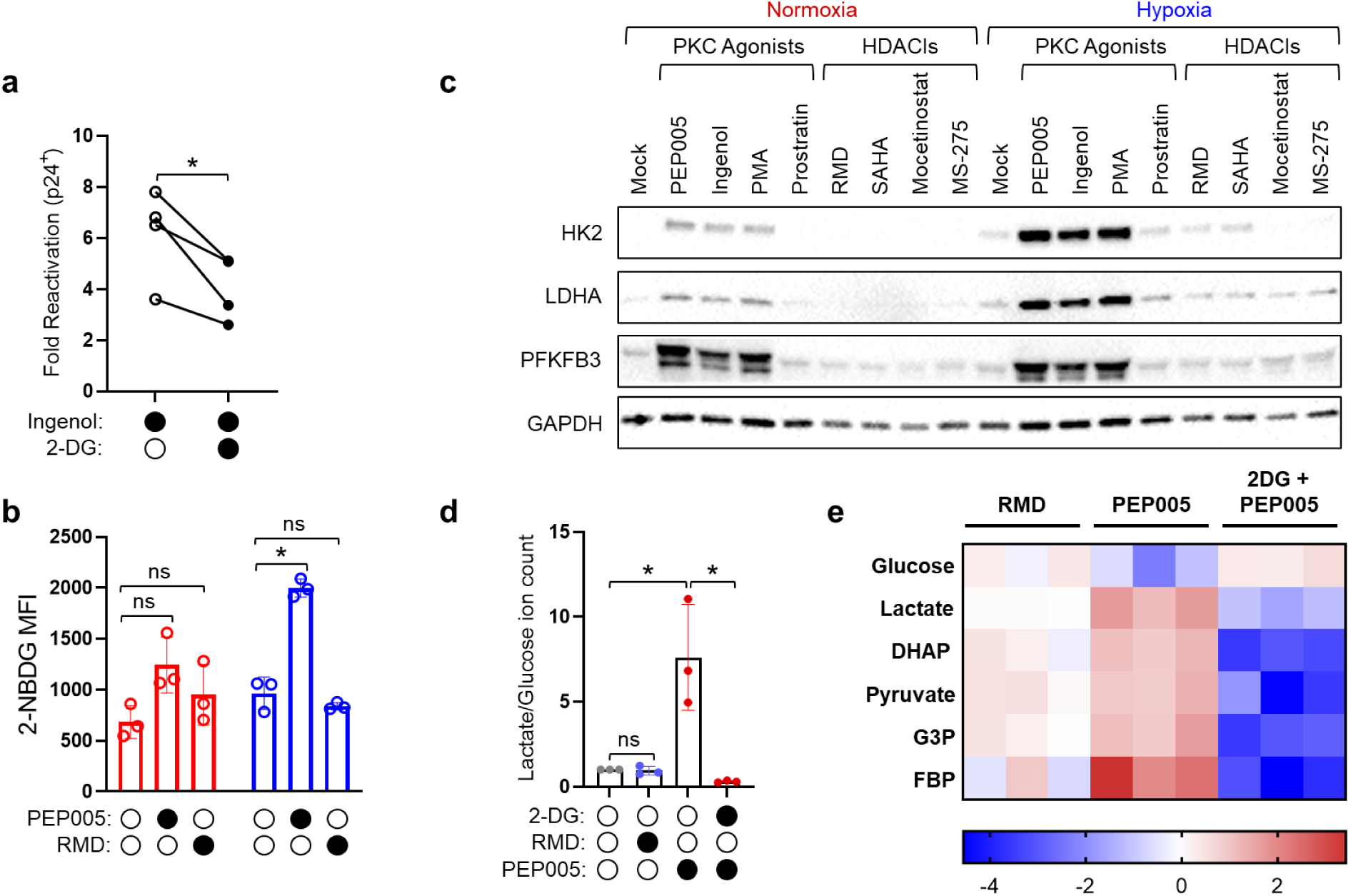
Latency reversal agents have differential capacities to induce glycolysis. **(a)** Reactivation of latent HIV-1 NL4-3 in an *in vitro* primary T_CM_ model of HIV latency established from 4 independent donor cells was measured by HIV p24 expression when latency reversal is stimulated by PKC agonist, ingenol-3,20-dibenzaote, in the absence or presence of glycolytic inhibitor, 2-deoxyglucose (2-DG, 5mM) as 2h pretreatment under normoxic (21% O_2_) conditions. Statistical significance was measured using paired t-test; *p<0.05. **(b)** Immunoblots of indicated glycolytic enzymes [hexokinase 2 (HK2), lactate dehydrogenase (LDHA) and fructose-2,6-biphosphatase 3 (PFKFB3)] assessed in whole-cell lysates of uninfected primary CD4 T cells either untreated or treated with a panel of PKC agonists [PEP005 (100nM), ingenol-3,20-dibenzoate (IDB, 100nM), PMA (100nM), prostratin (100nM)] or HDAC inhibitors [RMD (10nM), SAHA (2.5mM), mocetinostat (10mM), MS275 (10mM)] in 5mM glucose media for 48h under normoxic (21% O_2_) or hypoxic (1% O_2_) conditions. Glyceraldehyde 3-phosphate dehydrogenase (GAPDH) served as loading control. Immunoblots are representative of 3 replicates in independent donors. **(c)** Mean fluorescence intensity (MFI) of fluorescent glucose analog, 2-NBDG (400mM) in uninfected primary CD4 T cells either untreated or stimulated with PKC agonist, PEP005 (100nM) or HDAC inhibitor, RMD (10nM) for 48h under normoxic or hypoxic conditions to determine effects of LRAs on glucose uptake. Means and SDs of triplicate samples are shown. Statistical significance was determined using paired t-test, comparing each treatment to mock in each case; *p<0.05, ns = non-significant. **(d & e)** Global metabolomic analysis tracking the flux of **(d)** lactate and glucose ions and **(e)** indicated glycolytic intermediates using isotopic ^13^C in uninfected primary CD4 T cells that were either untreated or pretreated with glycolytic inhibitor, 2-deoxyglucose, 2-DG (5mM) for 2h under hypoxic (1% O_2_) conditions, then stimulated for 24h with PKC agonist, PEP005 (100nM) or HDAC inhibitor, RMD (10nM). Means and SDs of triplicate samples are shown. Statistical significance was measured using paired t-test; *p<0.05, ns = non-significant. In **(e),** heat map represents relative concentrations of each metabolite to untreated sample. Significant increases of p<0.05 are depicted in red and significant decreases of p<0.05 are depicted in blue.

To further determine the necessity of glucose in the reactivation of HIV, we assessed the ability of PEP005 and RMD to modulate glucose uptake. We demonstrated that while PEP005 significantly increases the uptake of glucose in T cells, RMD does not **(Fig. 2B)**. Next, to clearly demonstrate that these observations were not LRA-specific, we assessed the capacities of a panel of PKC agonists and HDAC inhibitors to induce expression of the tested glycolytic enzymes. Interestingly, all PKC agonists, except prostratin, despite being used at an equimolar concentration, could induce the expression of glycolytic rate-limiting enzymes hexokinase 2 (HK2), lactate dehydrogenase (LDHA), and 6- phosphofructo-2-kinase/fructose-2,6-biphosphatase 3 (PFKFB3) under both normoxia and hypoxia **(Fig. 2C)**. The inability of prostratin to induce expression of the glycolytic program could be linked to its much higher EC50 associated with inducing PKC downstream activity^45^. Consistent with the inability of RMD to induce glucose uptake in CD4 T cells, all HDAC inhibitors failed to induce the glycolytic program **(Fig. 2C)**. Induction of the glycolytic program by PKC agonists suggests they should increase glycolytic flux and lactate production in CD4 T cells. To test this directly, we performed a glucose tracing metabolomic analysis experiment. Here, as expected, PEP005, but not RMD increased glycolytic flux in treated CD4 T cells, as measured by increases in the ratio of labelled lactate ions to glucose ions **(Fig. 2D).** Consistent with this, we also observed increases in the relative abundance of radio-labeled lactate and other downstream glycolytic metabolites **(Fig. 2E)**. Importantly, like the sensitivity of HIV latency reversal to inhibition of glycolysis with 2-DG treatment, PEP005-stimulated increases in glycolytic flux was also suppressed by 2-DG treatment **(Fig. 2D, 2E)**. Collectively, our data suggests that LRAs from the PKC agonist class can increase the levels of lactate by inducing glycolytic flux via increasing the expression of glycolytic rate-limiting enzymes.

### Histone lysine lactylation in CD4 T cells is glycolysis-dependent and associated with HIV latency reversal

An established link exists between glucose-derived p ost-translational modifications like acetylation that promote HIV infection and latency reversal^25^. The role of acetylation of histones in the epigenetic regulation of HIV latency reversal have been widely studied, however, recent evidence suggests that other understudied acylations, including lactylation may play a role in proviral gene expression^50^. We therefore hypothesized that glucose-derived histone lactylation^42,51,52^ fueled by glycolysis may provide a mechanistic link between glucose availability and HIV latency reversal. To examine this potential molecular mechanism of glucose-induced reactivation of latent HIV, we assessed the capacity of LRAs to induce histone lactylation marks associated with increased gene transcription (H3K9la and H3K18la)^42,51,53–61^. To this end, we first performed immunoblotting of epigenetic marks at histone residue H3K9, a residue with modifications linked to HIV latency reversal and regulation of gene expression by metabolites and glucose-dependent lactylation^62,63^. Interestingly, we observed that PKC agonists that we previously found to induce glycolysis also increased histone H3K9 lactylation at both normoxia and hypoxia, but not H3K9 acetylation, a well-studied epigenetic mark associated with HIV reactivation^64^ **(Fig. 3A)**. By contrast, all tested HDAC inhibitors induced not only H3K9la, but also H3K9 acetylation without bias. Further, to confirm that LRA-mediated induction of H3K9la was glucose-dependent in our primary CD4 T cell system, we treated CD4 T cells with a representative PKC agonist (PEP005) in cell culture over a range of glucose concentrations. As expected, we found PEP005-mediated induction of H3K9la to be sensitive to glucose availability, with the modification suppressed at hypoglycemic conditions (0.625-2.5 mM glucose) **(Fig. 3B)**. Similarly, time-dependent PEP005-induced histone lactylation was potentiated by hypoxia and was also sensitive to glucose availability, as cells stimulated in the presence of 1mM glucose showed reduction in the accumulation of histone H3K9 lactylation (**Fig. S3**). Like the histone H3K9 residue, both acetylation and lactylation of the histone H3K18 residue are known to enhance gene expression^50,63,65^, but the dynamics of these two modifications at this residue have not been studied in the context of HIV latency reversal. To gain better insight into the potential role of induction of these two modifications at H3K18 in the mechanism of LRAs in HIV latency reversal, and since we show in **(Fig. 3A)** that HDAC inhibitors induce both modifications, we employed a HDACi LRA panel used in prior experiments **(Fig. 2B)** to assess how HIV reactivation was associated with the induction of H3K18lac **(Fig. 3C)** and H3K18ac **(Fig. 3D).** For this assessment, we utilized J-Lat 5A8 cells treated with titrated doses of each HDACi and determined how the level of both H3K18la and H3K18ac induced by each HDACi was associated with HIV reactivation stimulated by each compound. In these experiments we observed a greater association between reactivation and H3K18la (*r* = 0.6204, *P* = 0.0046) than with H3K18ac (*r* = 0.3939, *P* = ns). We also sought to determine the impact of HDACi treatment on the levels of H3K9ac and how well this modification was associated with HIV reactivation. As expected, we observed a significant association (*r* = 0.5158, *P* = 0.0238), although more modest than what we observed for H3K18la in treated cells **(Fig. 3E)**. Taken together, these data suggest that glucose-dependent histone lactylation could represent an additional biomarker for HIV latency reversal. This possibility is underscored by the fact that PKC agonists that potently induce HIV reactivation fail to induce histone acetylation at residues previously found to be associated with HIV latency reversal.

**Figure 3.**
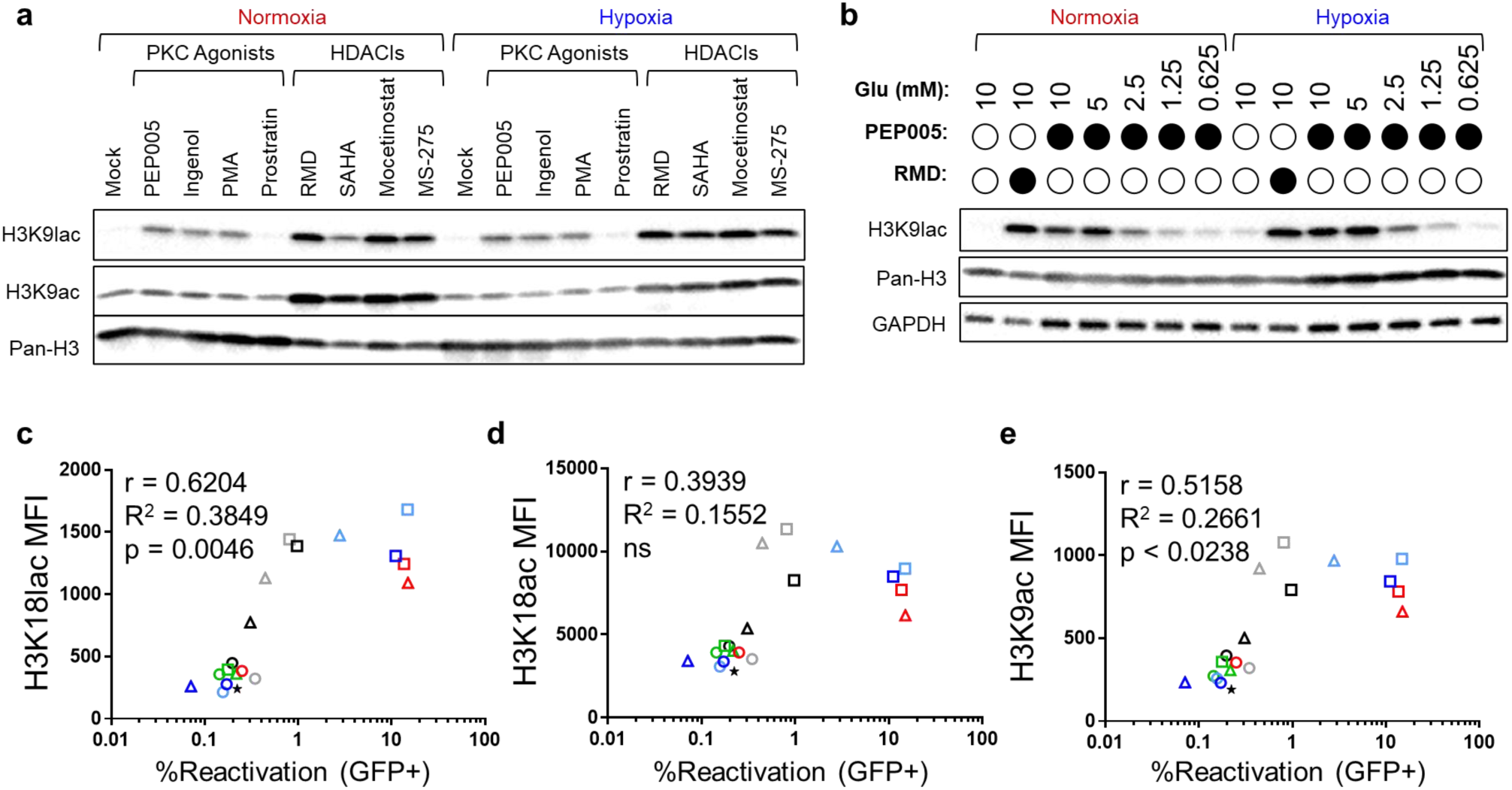
Histone lysine lactylation is induced by latency reversal agents, glucose dependent and associated with HIV latency reversal. **(a)** Immunoblots of histone 3 lysine 9 lactylation (H3K9lac) and acetylation (H3K9ac) assessed in whole-cell lysates of uninfected primary CD4 T cells either untreated or treated with a panel of PKC agonists [PEP005 (100nM), ingenol-3,20-dibenzoate (IDB, 100nM), PMA (100nM), prostratin (100nM)] or HDAC inhibitors [RMD (10nM), SAHA (2.5mM), mocetinostat (10mM), MS275 (10mM)] in 5mM glucose media for 48h under normoxic (21% O_2_) or hypoxic (1% O_2_) conditions. Total histone 3 served as loading control. Immunoblots are representative of 3 replicates in independent donors. **(b)** Immunoblot of H3K9lac assessed in whole-cell lysates of uninfected primary CD4 T cells either untreated or treated with HDAC inhibitor, RMD in 10mM glucose media or PKC agonist, PEP005 over a range of glucose concentrations as indicated for 48h under normoxic (21% O_2_) or hypoxic (1% O_2_) conditions. Total histone 3 and GAPDH served as loading control. Immunoblots are representative of 3 replicates in independent donors. **(c - e).** Correlation analyses of HIV reactivation measured by GFP positivity with post-translational modifications [**(c)** H3K18lac, **(d)** H3K18ac and **(e)** H3K9ac] in J-Lat 5A8 cells that were untreated or treated with titrated HDAC inhibitors for 24h under hypoxic (1% O_2_) conditions as follows: RMD (10nM), BUT (0.5mM), mocetinostat (1, 0.2, 0.04μM), MS275 (1, 0.2, 0.04μM), RGFP966 (1, 0.2, 0.04μM), SAHA (1, 0.2, 0.04μM) and trichostatin (TSA, 1, 0.2, 0.04μM). p>0.05 = ns (non-significant).

Given the impact of glucose availability on histone marks that drive gene expression and HIV latency reversal, and the fact that histone lactyl-transferases EP300 and KAT2A have both been implicated in the regulation of HIV latency, we hypothesized that these epigenetic writers link glucose-dependent lactate production to histone-modifications that promote HIV latency reversal^66,67^. Thus, we investigated whether these epigenetic writers were mediating the effects of LRAs in HIV reactivation via modulating histone lactylation marks. First, to formally test the role of both EP300 and KAT2A in latency reversal in our system, we treated J-LAT 5A8 cells with A-485 and MB-3, specific small molecule inhibitors of EP300 and KAT2A, respectively **(Fig. 4A-B)**. A-485 treatment did not inhibit PEP005-induced latency reversal **(Fig. 4A)**. By contrast, MB-3 suppressed PEP005- mediated latency reversal under both hypoxia and normoxia, but to a lesser extent under the latter **(Fig. 4B)**. Next, we directly examined the impact of EP300 and KAT2A inhibition on glucose-dependent histone modifications catalyzed by these lactyl-transferases in CD4 T cells. We observed that inhibition of KAT2A with MB-3 reduced both lactylation and acetylation of H3K9 in a concentration-dependent manner under hypoxic conditions **(Fig. 4C)**. Inhibition of EP300 with A-485 did not appreciably impact either of these modifications, but did reduce acetylation of histone H3K27, a well-documented EP300 target. These results are consistent with a role of KAT2A in mediating PEP005-dependent latency reversal via catalyzing glucose-dependent histone modifications.

**Figure 4.**
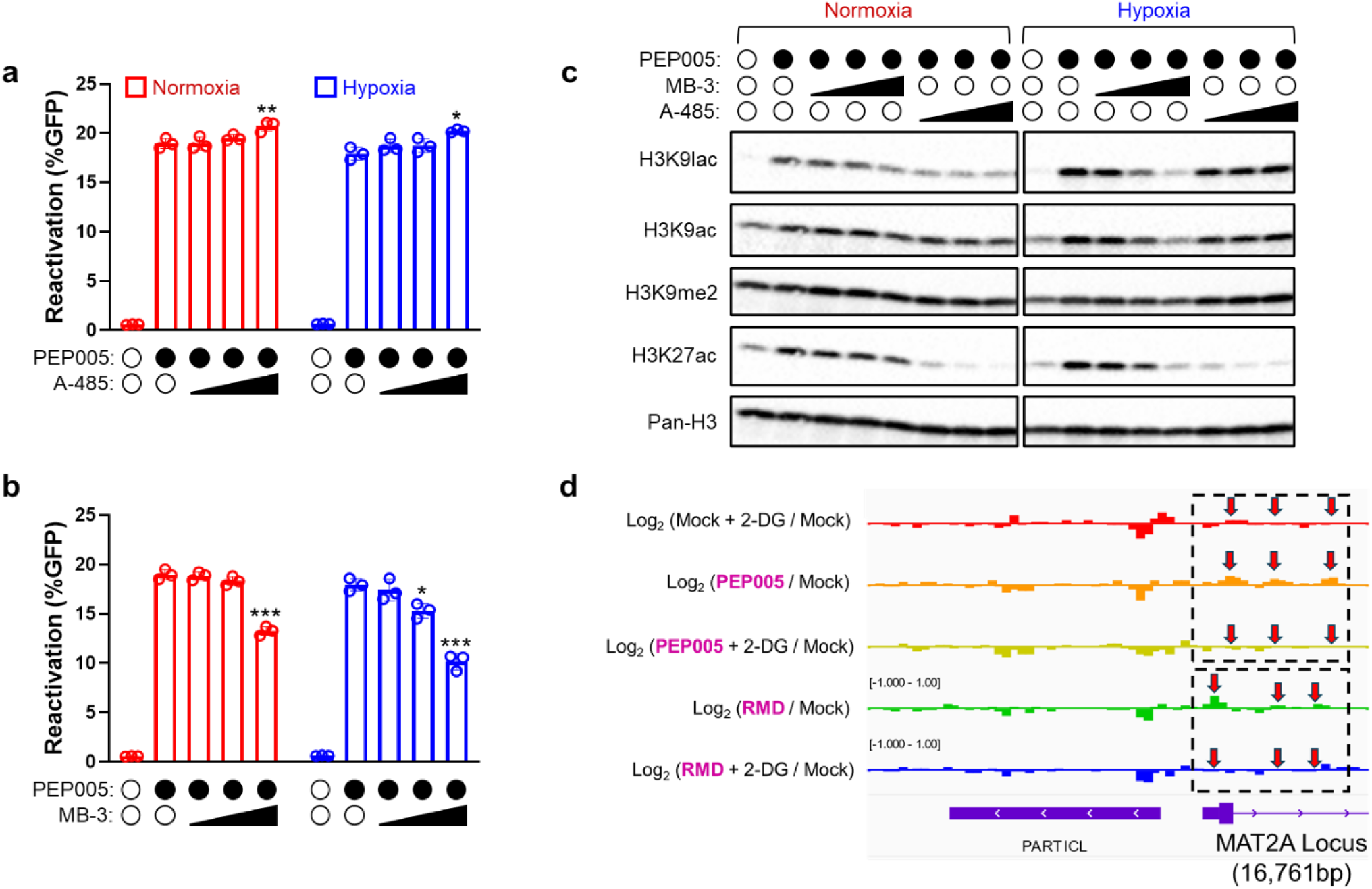
Activity of histone acyl-ltransferase, KAT2A is associated with histone lactylation and HIV latency reversal. **(a & b).** Reactivation of latent HIV in J-Lat 5A8 cells measured by GFP fluorescence when cells are either untreated or stimulated for 24h with PKC agonist, PEP005 (100nM) in the absence or presence of titrated **(a)** KAT2A inhibitor, butyrolactone-3 (MB-3) at 25, 50 and 100uM and **(b)** p300/CBP inhibitor, A-485 at 2.5, 5 and 10uM under normoxic (21% O_2_) or hypoxic (1% O_2_) conditions. Means and SDs of triplicate samples are shown. Statistical significance was determined using paired t-test, comparing reactivation in the presence of inhibitors to reactivation in the absence of inhibitors; *p<0.05, **p<0.005, ***p<0.0005. **(c)** Immunoblots of post-translational modifications [H3K9lac, H3K9ac, histone 3 lysine 9 demethylation (H3K9me2) and histone 3 lysine 27] acetylation assessed in whole-cell lysates of uninfected primary CD4 T cells either untreated or treated with 25, 50 and 100uM MB-3 or 2.5, 5 and 10uM under normoxic (21% O_2_) or hypoxic (1% O_2_) conditions. Total histone 3 served as loading control. Immunoblots are representative of 3 replicates in independent donors. **(d). Glycolytic inhibition limits proviral chromatin accessibility.** ATAC-seq analysis of proviral chromatin accessibility in J-Lat 5A8 cells that were either untreated or pretreated with glycolytic inhibitor, 2-deoxyglucose, 2-DG (5mM) for 2h under hypoxic (1% O_2_) conditions, then stimulated for 24h with PKC agonist, PEP005 (100nM) or HDAC inhibitor, RMD (10nM). Normalized sequencing tracks for the respective treatments are presented for the MAT2A locus (indicated with red arrows) which contains the integrated provirus in the J-Lat 5A8 clone (coordinate chr2:85,758,801-85,775,562; 16,761bp). Average accessibility of 3 replicates are shown for each treatment.

### Glycolysis inhibition corresponds to loss of chromatin accessibility at the HIV LTR

Glucose-dependent histone acetylation and lactylation modifications are associated with increased gene expression across systems^42,51,52,59,68–74^. We proposed that glycolysis, via these modifications, promotes chromatin accessibility to facilitate proviral gene expression. To this end, we performed ATAC-seq and accessed the MAT2A locus which is the integration site of the provirus in the J-Lat 5A8 cells. In line with our hypothesis, we observed that PKC agonist PEP005 and HDACi RMD both increased chromatin accessibility relative to mock-treated cells at the HIV LTR. To directly test the role of the metabolism of glucose in facilitating the observed increase in chromatin accessibility, we treated cells with 2-DG to block glycolytic flux. As predicted, we showed that inhibition of glycolysis suppressed the increases in chromatin accessibility induced with either PEP005 or RMD **(Fig. 4D)**. This is consistent with the requirement of glucose to fuel the histone marks enhanced by both treatments. This demonstrates a mechanistic link between glucose metabolism and chromatin accessibility necessary for HIV latency reversal.

## Discussion

The barrier to a functional cure for HIV is the latent proviral reservoir which remains refractory to ART. Significant efforts have been made to purge this latent reservoir with LRAs identified in vitro, but these agents were found to be ineffective in clinical trials^75^. An increased understanding of the determinants of latency reversal could facilitate the development of strategies that effectively target the HIV reservoir in vivo. To this end, we sought to define the role of the central metabolic pathway, glycolysis, in latency reversal vis-a-vis prior associations with both virus replication during primary infection^25,26,34,76^ and virus rebound from latency^27,29,36^. A major consideration in our approach to this study was modeling physiologic hypoxia, a key feature that has a profound impact on the metabolism of CD4 T cells and tissues that harbor the HIV reservoir *in vivo*^22^. Using multiple complimentary approaches, we showed that glycolysis is a cellular determinant that impacts latency reversal, especially under hypoxic conditions (1% O_2_). This observation is of particular importance since most HIV latency reversal studies were executed *in vitro* using standard tissue culture conditions at normoxia. Importantly, the metabolic capacity of cells to utilize the TCA cycle is markedly compromised in hypoxic microenvironments^77^, thereby increasing cellular dependence on aerobic glycolysis as the primary source of energy, co-factors, and epigenetic metabolites needed to regulate gene expression. This contrasts with normoxic microenvironments where cells exhibit metabolic flexibility and undergo metabolic rewiring to utilize the TCA cycle to compensate for when glycolysis is compromised.

The current study also demonstrates that the intrinsic capacity of LRAs, modeled by PKC agonists, to induce glycolytic flux to fuel latency reversal at physiological levels of oxygen and glucose, thereby preserving LRA efficacy under conditions which characterize target tissue reservoirs *in vivo*. We clearly show that “nutrient-rich” conditions with supranormal levels of atmospheric oxygen levels and glucose, as modeled by standard tissue culture conditions, mask the true metabolic dependencies of LRAs during *in vitro* studies. We posit that the inability of HDAC inhibitors to induce glycolytic flux may be one of several reasons why they fail to consistently elicit productive reactivation (viremia) or reduce reservoir size *in vivo*^78–81^ despite encouraging *in vitro* efficacies in pre-clinical trials^82^. These findings interestingly highlight the induction of glycolysis as another key functionality of the PKC agonist class of LRAs which hitherto have principally been associated with mediating latency reversal via NF-κB activation^83^.

Glycolysis is known to fuel histone lactylation even at hypoxia^42,56^. We have identified KAT2A as the critical lactyl-transferase that transfers the lactyl group from glucose and lactate-derived lactyl-CoA to histone H3 lactylation^67^. Notably, histone H3 is preferentially lactylated rather than acetylated even in the presence of equimolar lactyl-CoA and acetyl-CoA, as the Km of KAT2A for lactyl-CoA is lower than that for acetyl-CoA^67^. We surmise that hypoxia, a condition which increases glycolysis and lactate production, facilitates HIV latency reversal via increased histone H3 lactylation, rather than acetylation, because hypoxia further reduces the available pool of the acetyl-CoA substrate, thus enhancing KAT2A lactyl-transferase activity^67^.

Additionally, we found latency reversal by PKC agonists was not associated with an increase in acetylation at histone H3K9, a principal modification necessary for displacement of repressive complexes during latency reversal^64,84,85^. Rather, PKC agonists selectively increased global histone lactylation^42,86–89^, while HDAC inhibitors that directly inhibit class I HDACs (which are also histone delactylases)^90^ increased both modifications at H3K9. Recent studies have suggested that latency prevention by class I HDAC inhibitors is associated with increased H3K9 acetylation and this epigenetic modification was a key determinant of HIV latency^64^. Our results indicate that H3K9 lactylation may also play a role in HIV latency, as we found PKC agonists failed to induce global histone acetylation, suggesting that acetylation at histone H3K9 was not necessary for PKC agonist-mediated latency reversal. Thus, this observation may explain why the induction of histone H3 acetylation by HDAC inhibitors *in vivo* does not strongly corelate with virus reactivation^62,91^. Furthermore, to the best of our knowledge, this is the first study that demonstrates an association between histone lactylation and HIV latency reversal.

In sum, our study highlights glycolysis as a major driver of HIV latency reversal that provides insight into the links between aerobic glycolysis, its end products, and virus rebound in PLWH. Mechanistically, the current study connects glycolysis to histone lactylation, a post-translational modification not hitherto linked to HIV biology. This links our work to previous studies that have defined an indispensable role for mTOR-dependent processes in HIV latency reversal, as mTOR represents a central regulator of glycolysis in T cells^92^. Our findings suggest that the ability of LRAs to induce glycolysis may favor their utility *in vivo*, and that lactylation may be a promising biomarker for latency reversal *in vivo* and for pharmacological screenings of compounds moving into clinical trials. Lastly, we show that modeling relevant physiological conditions such as hypoxia is indispensable for uncovering the metabolic dependencies of intricate processes such as latency reversal, informing experimental approaches in the field of HIV research.

## Materials and Methods

### Cell culture

J-Lat cells (The 5A8 clone was kindly provided by Dr. Warner Greene, Gladstone Institute, UCSF) or primary human CD4 T cells were cultured in Roswell Park Memorial Institute (RPMI)-1640 (Corning) supplemented with 10% Fetal Bovine Serum (FBS) (Cytiva) and 1% penicillin/streptomycin (Gibco) and maintained at 37°C with 21% oxygen and 5% carbon dioxide. For hypoxic experimentation, cells were maintained in a specialized hypoxic workstation (Whitley H35 HEPA hypoxystation) at 37°C with 1% oxygen, 5% carbon dioxide and 94% nitrogen for the indicated time in each experiment.

### Latency reactivation assays in J-Lat cells

J-Lat cells were plated at 0.5 x 10^6^ cells/ml/well in 24 well plates. Identical sets of plates were incubated under normoxic (21% O_2_) versus hypoxic (1% O_2_) conditions for all experiments. In the standard latency reversal assays, cells were pretreated for 2h with inhibitors: 2-deoxyglucose (Cayman Chemical), sodium oxamate (Cayman Chemical), CB-839 (Selleckchem), BMS345541 (Selleckchem), AZD2014 (Cayman Chemical) or A485 (Cayman Chemical) as indicated, and reactivated from latency with PKC agonists [ingenol-3-angelate, PEP005 (Cayman Chemical), ingenol-3, 20-dibenzoate, IDB (Enzo Life Sciences), PMA (Sigma), byrostatin (Sigma), prostratin (Tocris Bioscience)], HDAC inhibitors [romidepsin, RMD (Cayman Chemical), SAHA (Cayman Chemical), sodium butyrate (Sigma-Aldrich), MS275 (Cayman Chemical), trichostatin A (Cayman), RGFP966 (Cayman Chemical), mocetinostat (Cayman Chemical)] or a combination of PEP005 and romidepsin in the synergy experiments. In add-back experiments, sodium L-lactate (Sigma-Aldrich), sodium pyruvate (Sigma-Aldrich) or D-glucose (Gibco) was added at the time of stimulation with LRAs. Cells were harvested after 24h and analyzed by flow cytometry or western blots. In the metabolite substitution experiments, cells were washed once in glucose-free media and then plated in glucose-free media supplemented with matched concentrations of galactose (Sigma) for 24h.

### Primary human resting CD4 T cell isolation

Isolation of human CD4 T cells from PBMCs was done as previously described^93^. Briefly, PBMCs were isolated from the blood of healthy, deidentified donors (New York Blood Center) by centrifuging the blood through a Ficoll-Paque Plus (GE Healthcare) gradient to separate plasma and red blood cells from the PBMCs. PBMCs were then washed repeatedly with phosphate buffered saline, and resting CD4+ T cells were isolated from negatively-selected total CD4 T cells (CD4 T cell isolation kit (human), Miltenyi Biotec) using CD25 and HLA-DR microbeads (Miltenyi Biotec) according to the manufacturer’s instructions. In general, primary T cell experiments involved 2h inhibitor pretreatments and LRA treatment for 48h before harvest.

### *In vitro* primary T_CM_ cell model of latency

Establishment of a primary T_CM_ latency model was as previously described^46^ in the laboratory of Dr Alberto Bosque (George Washington University). Briefly, naïve CD4 T cells were activated with anti-CD3/anti- CD28 Dynabeads (1:1 ratio of cells to beads) in the presence of anti-human IL-4 (1 μg/ml, Peprotech), anti-human IL-12 (2 μg/ml, Peprotech), and TGF-β1 (10 ng/ml, Peprotech). Cells were plated in 96-well round-bottom plates at a density of 0.5 × 10^6^/ml in RPMI supplemented with 10% FBS, penicillin/streptomycin, and L-glutamine (complete RPMI medium) for 3 days. Afterwards, anti-CD3/anti-CD28 beads were removed magnetically using a Dynal MPC-L magnetic particle concentrator (Invitrogen). Cells were resuspended and maintained at a density of 1 × 10^6^/ml in complete RPMI medium with 30 IU/ml of IL-2. Medium was replaced on days 4 and 5 of culture. To establish latency, one-fifth of cells were infected on day 7 of culture with NL4-3 virus by spinoculation (2,900rpm, 2h, 37°C) while one-fifth served as uninfected control. After spinoculation, infected cells were added to the remaining three-fifths of culture with complete medium and IL-2. At day 10, cells were plated in 96-well round-bottom plates in complete medium with 30 IU/ml IL-2 to facilitate cell-to-cell spread of infection (“crowding” phase). At day 13, cells were transferred to flasks and the following antiretroviral drugs were added to both infected and uninfected cultures to stop further infection: 0.5 μM nelfinavir, 100 nM efavirenz, and 100 nM AMD-3100^46^. On day 17, infected and uninfected cells were sorted using a CD4 positive isolation kit (Dynabeads, 11331D; Invitrogen) to isolate the latently infected cell population. The isolation was carried out as indicated in the manufacturer’s protocol, with two modifications: (1) the amount of CD4 beads was increased 3-fold and (2) the resuspension volume of buffer II was changed to 200 to 300 μl per 10^7^ cells. For the reactivation assay, latently infected cells were either mock or pretreated with 2-DG for 2h and reactivated with ingenol-3,20-dibenzoate or anti- CD3/anti-CD38 beads for 48h before flow cytometry analyses.

### Flow cytometry analyses and antibodies

To exclude the non-viable cells in standard latency reactivation assays, cell pellets were resuspended in cold SYTOX blue dead cell stain (Thermo Fisher Scientific) [in FACS buffer (1x PBS/2mM EDTA/1% BSA)] and immediately analyzed by flow cytometry. For intracellular staining for HIV p24 or post-translational modifications (EpiFLOW), cells were harvested and stained for viability with aqua fluorescent reactive dye (Thermo Fischer Scientific) for 30mins at 4°C, washed once in ice cold FACS buffer, then subsequently fixed and permeabilized with Cytofix/Cytoperm solution (BD Biosciences). The permeabilized cells are then washed with perm/wash (BD Biosciences) and stained with the following antibodies: H3K9ac-PacBlue (Cell Signaling Technology, #11857S), H3K18ac-A647 (Cell Signaling Technology, 20363S), H3K18ac- AF488 (Cell Signaling Technology, 73508S), H3K18lac-PE (PTM) or p24-APC (MediMabs). The H3K18lac antibody was conjugated to PE using a SiteClick Antibody Labeling Kit (R-Phycoerythrin) (Thermo Fisher Scientific) according to manufacturer’s instructions. Thereafter, cells were diluted in perm/wash buffer and analyzed by flow cytometry. Flow cytometric data was obtained on a LSRFortessa (BD) and analyzed using FlowJo software.

### Western blots

Cell pellets were lysed for 30mins on ice using Pierce^TM^ RIPA buffer (Thermo Scientific) supplemented with protease inhibitor (Thermo), phosphatase inhibitor (Roche) and universal nuclease (Thermo Scientific). Clarified lysates (centrifuged at 16,000 x g for 20mins) solubilized in laemmli sample buffer (Bio-Rad) and dithiothreitol (DTT; Sigma) were denatured at 95°C for 5mins, resolved by SDS-PAGE on 4 - 15% Mini-PROTEAN TGX gels (Bio-Rad) and transferred to a nitrocellulose membrane. Blots were blocked in TBS Superblock^TM^ Blocking Buffer (Thermo Scientific) for 1h at room temperature, and incubated with indicated primary antibodies overnight on a rocker at 4°C. Following sequential washes with TBST (1X TBS + 0.1% Tween20), the membranes were incubated for 1h at room temperature with HRP-linked secondary antibodies diluted in TBST supplemented with 0.5% w/v Blotto (Santa Cruz Biotechnology). After repeated washes, the immunoreactive protein bands were visualized by Chemiluminescence using a ChemiDoc XRS+ Gel Imaging System (Bio-Rad) according to the manufacturer’s instructions. Image analysis was performed using Image Lab software. The following antibodies were used: HIF-1α (Proteintech, #20960-1-AP), Phospho GLUT-1 S226 (Sigma-Aldrich, #ABN991), GLUT-1 (EMD Millipore Corp, #07-1401), GLUT-3 (Proteintech, #20403-1-AP), HK2 (Cell Signaling Technology, #2867T), PFKFB3 (BETHYL, #A304-249A-T), LDHA (Cell Signaling Technology; #3582S), Phospho S6 S240/244 (Cell Signaling Technology; #5364T), S6 (Cell Signaling Technology; #2217S), Phospho NF-κB p65 (Ser536) (Cell Signaling Technology, #3033), NF-κB p65 (Santa Cruz, #sc-372) H3K9ac (Cell Signaling Technology, #9649T), H3K27ac (Cell Signaling Technology; #8173T), pan lactyl lysine (PTM; #1401), H3K9lac (PTM; #1419RM), H4K8lac (PTM; #1415), pan H3 (Cell Signaling Technology; #4499T) and histone H3 (96C10) (cell Signaling Technology, #3638), GAPDH-HRP (Cell Signaling Technology; #8884S) and anti-rabbit IgG-HRP (Cell Signaling Technology, #7074S and Thermo Fischer Invitrogen, #G21234).

### Glucose uptake assay

Primary CD4 T cells were plated and treated with LRAs as indicated. At the 48h harvest time point, cells were harvested, pelleted (300 x g, 5mins) and washed in 1ml of pre-warmed glucose free RPMI. Cells were then resuspended in the glucose free RPMI (200ul), incubated for 30 min at 37°C, then treated with fluorescent glucose analog, 2-NBDG (400 mM, GLPBIO) for 15 min at 37°C, shielded from light, before washing, viability staining and flow cytometry analysis.

### Isotopic glucose tracing

Water soluble metabolites: Primary CD4 T cells were isotopically labeled with [U-^13^C] Glucose (Cambridge Isotope Laboratories) starting during a 2h pretreatment with 2-DG and stimulation with PEP or RMD as indicated. Cell pellets were extracted in 80% Methanol and vortexed for 25 seconds and incubated on dry ice for 10 minutes. Cell samples were then centrifuged at 16,000 g for 30 minutes. The supernatants were transferred to new Eppendorf tubes and then centrifuged again at 16,000 g for 25 minutes to remove and residual debris before analysis. The supernatants were centrifuged at 16,000 g for 20 minutes to remove any residual debris before analysis. Supernatants were analyzed within 24 hours by liquid chromatography coupled to a mass spectrometer (LC-MS).

Extracts were analyzed within 24 hours by liquid chromatography coupled to a mass spectrometer (LC-MS). The LC–MS method was based on hydrophilic interaction chromatography (HILIC) coupled to the Orbitrap Exploris 240 mass spectrometer (Thermo Scientific)^94^. The LC separation was performed on a XBridge BEH Amide column (2.1 x 150 mm, 3.5 𝜇m particle size, Waters, Milford, MA). Solvent A is 95%: 5% H2O: acetonitrile with 20 mM ammonium acetate and 20mM ammonium hydroxide, and solvent B is 90%: 10% acetonitrile: H2O with 20 mM ammonium acetate and 20mM ammonium hydroxide. The gradient was 0 min, 90% B; 2 min, 90% B; 3 min, 75% B; 5 min, 75% B; 6 min, 75% B; 7 min, 75% B; 8 min, 70% B; 9 min, 70% B; 10 min, 50% B; 12 min, 50% B; 13 min, 25% B; 14min, 25% B; 16 min, 0% B; 18 min, 0% B; 20 min, 0% B; 21 min, 90% B; 25 min, 90% B. The following parameters were maintained during the LC analysis: flow rate 150 mL/min, column temperature 25 °C, injection volume 5 µL and autosampler temperature was 5°C. For the detection of metabolites, the mass spectrometer was operated in both negative and positive ion mode. The following parameters were maintained during the MS analysis: resolution of 180,000 at m/z 200, automatic gain control (AGC) target at 3e6, maximum injection time of 30 ms and scan range of m/z 70-1000. Raw LC/MS data were converted to mzXML format using the command line “msconvert” utility^95^. Data were analyzed via the MAVEN software, and all isotope labeling patterns were corrected for natural ^13^C abundance using AccuCor^96^.

### ATAC-seq

ATAC-seq was performed at the Biotechnology Resource Center (BRC) Epigenomics Facility (RRID:SCR_021287) at the Cornell Institute of Biotechnology. Briefly, 50,000 J-Lat cells flash-frozen in cryopreservative (10% DMSO (Sigma-Aldrich) in FBS) were lysed, permeabilized, and tagmented using the Omni ATAC-seq protocol^97^ followed by 12 cycles of barcoding PCR. Gel-purified libraries were sequenced on the NextSeq500 (CU BRC Genomics Facility; RRID:SCR_021727) to obtain 10M paired-end reads.

## Acknowledgements

The authors thank William Lai and Amy Lyndaker at the Biotechnology Resource Center (BRC) Epigenomics Facility (RRID:SCR_021287) at the Cornell Institute of Biotechnology for ATAC-seq experiments. We would also like to acknowledge the Huck Institutes’ Metabolomics Core Facility (RRID:SCR_023864) for use of the OE 240 LCMS and Sergei Koshkin for helpful discussions on sample preparation and analysis. The authors are also grateful for the technical contributions of Nicole M. Andre and Jocelyn Kim.

This work was supported by NIH grant U54 AI170660 (to HET), The Howard Hughes Medical Institute Hanna H. Gray Fellows Program Faculty Phase (Grant# GT15655 to MRM) and the Burroughs Welcome Fund PDEP Transition to Faculty (Grant# 1022604 to MRM). GES was supported by NCI Career Development Award K01CA255406. EKM was supported by NIH NIAID-funded T32 Grant (T32CA247756). This research has been facilitated by the services and resources provided by District of Columbia, Center for AIDS research, an NIH funded program AI117970 (to AB), which is supported by the following NIH co-funding and participating institutes and centers: NIAID, NCI, NICJD, NHLBI, NIDA, NINH, NIA, FIC, NIGGIS and NDDK and OAR.

---

**Figure S1.**
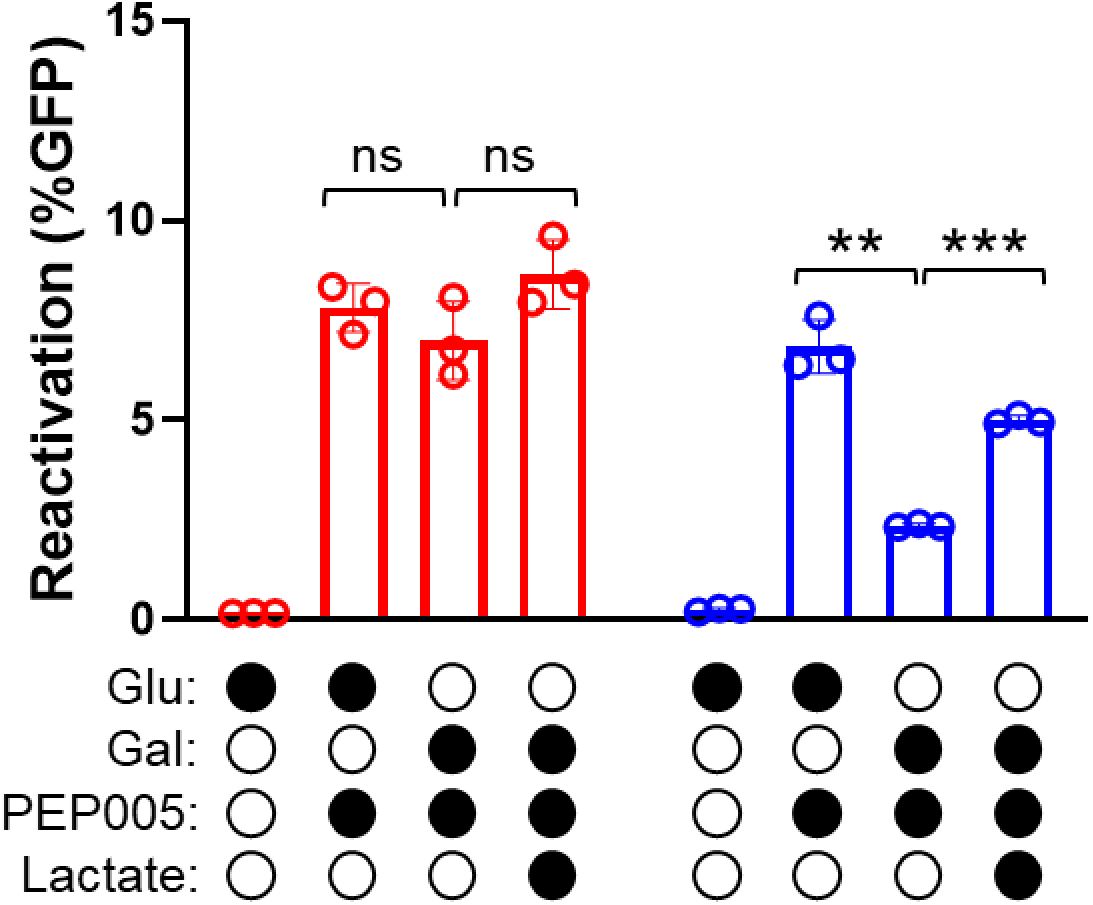
Metabolite substitution demonstrates the dependence of HIV latency reversal on glucose metabolism. Reactivation of HIV latency assessed by GFP expression in J-Lat 5A8 cells that were either unstimulated or stimulated for 24h with PKC agonist, PEP005 (100nM) in either 5.5mM glucose- or 5.5mM galactose-containing media under normoxia (21% O_2_) or hypoxia (1% O_2_) with or without supplementation with sodium lactate (50mM) in galactose medium. Means and SDs of triplicate samples are shown. Statistical significance was determined using paired t-test; **p<0.005, ***p<0.0005, ns = non-significant.

**Figure S2.**
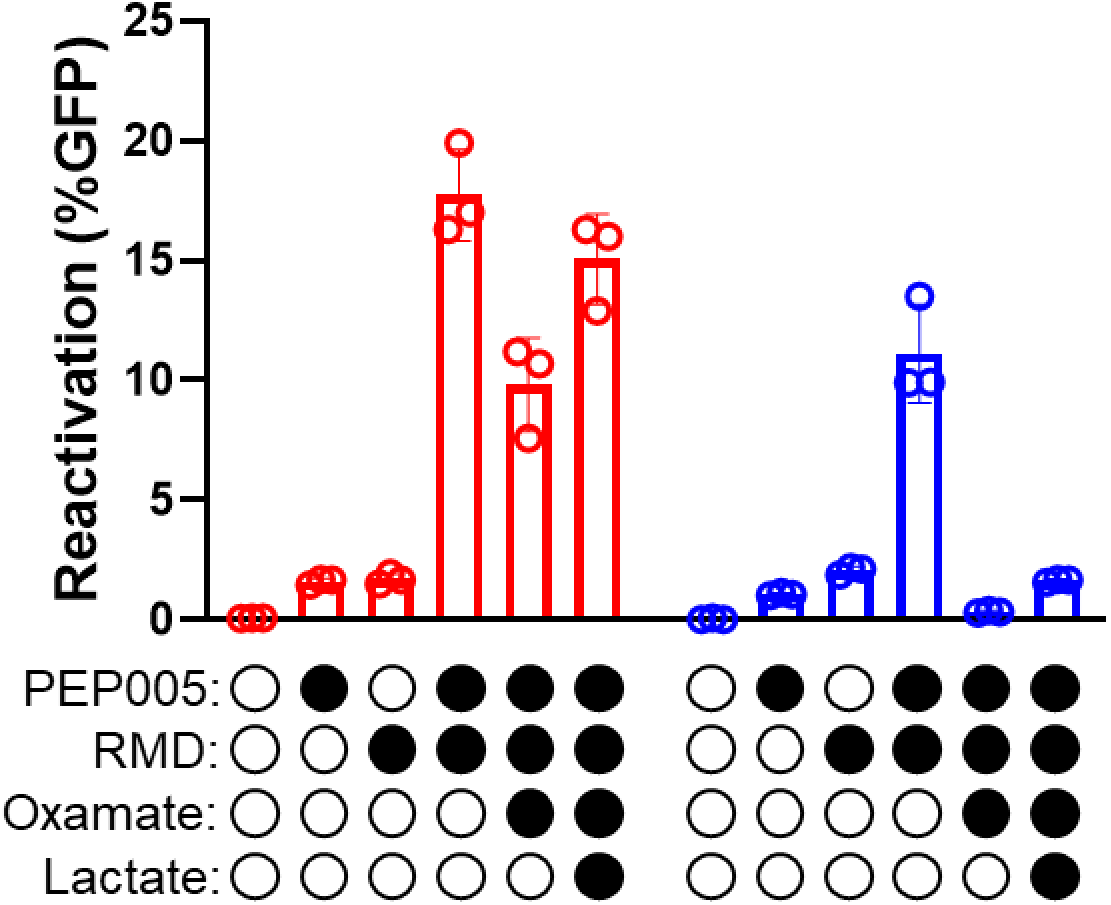
HIV latency reversal by combinatorial LRA treatment is dependent on glycolysis. Reactivation of HIV latency assessed by GFP expression in J-Lat 5A8 cells either unstimulated or stimulated for 24h with PKC agonist, PEP005 (10nM) or HDAC inhibitor, RMD (4nM) or their combination, with or without pretreatment with glycolytic inhibitor, sodium oxamate (25mM) under normoxic (21% O_2_) or hypoxic (1% O_2_) conditions. Inhibitor treatment was with or without supplementation with exogenous sodium lactate (50mM) during stimulation. Means and SDs of triplicate samples are shown.

**Figure S3.**
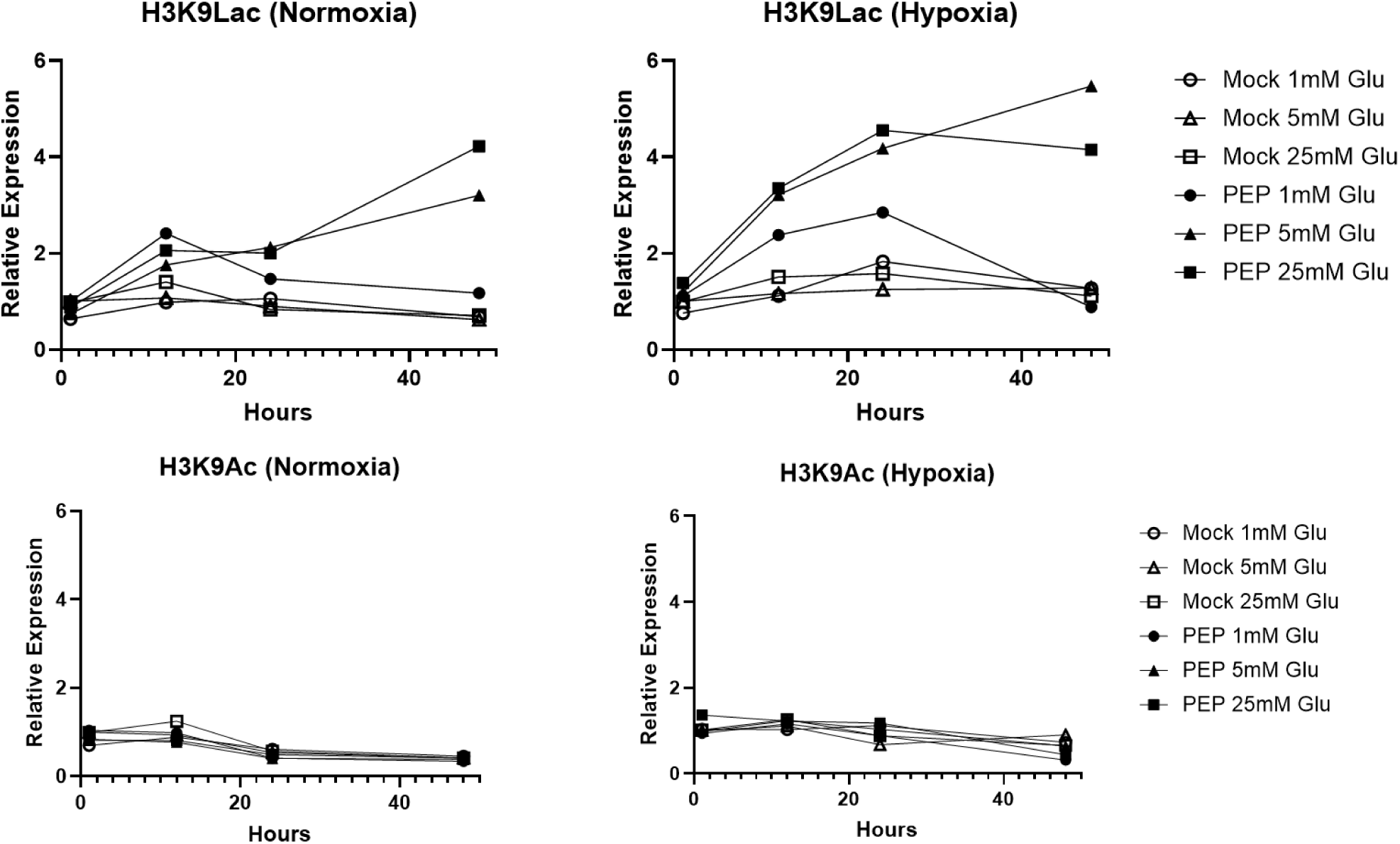
Time course experiment reveals the glycolytic dependence of histone lysine lactylation in a temporal fashion. **(a-d).** Graphs show the relative fold change of densitometry values from a representative immunoblot of H3K9lac or H3K9ac induced under normoxic (21% O_2_) or hypoxic (1% O_2_) conditions in uninfected primary CD4+ T cells either unstimulated or stimulated with PKC agonist, PEP005 in media containing 1, 5 or 25mM glucose for 1, 12, 24 or 48h. 21% O_2_) or hypoxia (1% O_2_) respectively. Densitometry analyses is representative of data from 3 replicates in independent donors and was performed with Image Lab software.

